# Ancient occupation by humans leads to missing bird diversity in otherwise natural habitats

**DOI:** 10.1101/2024.01.23.576512

**Authors:** Alexander H. Murray, Luke O. Frishkoff

## Abstract

Habitat modification is responsible for substantial biodiversity declines, but communities vary in their tolerance to land-use change. One infrequently queried possibility is that historical factors determine the sensitivity of contemporary communities. We use bird community data from 54 studies across the world to test the hypothesis that pre-historic human presence reduced community sensitivity to land-use change by eliminating sensitive species in natural habitats. We find that pre-historic human population size correlates with reduced sensitivity of communities. Primary vegetation in areas with larger pre-historic human populations contain fewer species today, while species richness in structurally simple agriculture is unimpacted. The greatest signal of humans impacts dates back to 12,000 YBP suggesting that early humans may have caused even more widespread extinctions, than previously appreciated.

**One-Sentence Summary:** Areas with high human population 12,000 years ago have less biodiversity today, but are more tolerant of habitat modification

## Main Text

Biodiversity is declining, and an imperative for the 21^st^ century is to understand why. Ongoing anthropogenic influences such as habitat modification, climate change and invasive species contribute to many declines (*1*). Amongst these, habitat modification is responsible for threatening the greatest number of species but the degree to which habitat modification results in declines varies substantially between regions and species. Modern day conditions account for some of this variation, as both the severity of land use change (*2-4*) and climatological context alter outcomes of land-use change (*3*). For example, the effects of habitat loss and fragmentation are most negative in areas with warm maximum temperatures (*5*). But an intriguing possibility is that historical factors also play a key role in explaining modern variation in biodiversity loss due to habitat conversion. The extinction filter hypothesis (*6*) proposes such a phenomena, in which a biological community’s prior exposure to disturbance has eliminated sensitive species, thereby predisposing the community as a whole to appear highly resistant to the same disturbance in the future. If this were the case, then areas with long histories of human disturbance would have lost many species sensitive to disturbance, rendering the modern day assemblage resistant to biodiversity loss due to land-use change. But an alternative mechanism could generate a similar pattern of resilience. Some species exposed to human disturbance find ways of coping (e.g. through evolutionarily adaptation). If such adaptation is widespread, areas with long-standing human influence would possess assemblages that are more resilient to human impacts.

Support for the extinction filter hypothesis exists for some natural forms of habitat change. Communities which have been subjected to more frequent natural disturbances tend to contain species that are less sensitive to human-caused forest loss (*7*). Regions impacted by historical disturbance (fire, storms and deforestation) have fewer forest-dependent species remaining today (*8*), and conversely, in the most intact landscapes new habitat modification has the most damaging impacts for biodiversity (*9*). Further, archeological evidence supports the idea that (pre-)historical human impacts may have had large biodiversity consequences. Wherever humans appeared as they spread out of Africa, extinctions followed in short time (*10*). In Australia human-induced extinctions may date back 40-50,000 YBP (years before present) when large mammals were lost shortly after human arrival on the continent (*11, 12*). Similar declines occurred in in the Americas when extinctions of megafauna coincided with human arrival and spread (*13, 14*). While the megafauna extinctions are most prominently documented, prehistoric extinctions caused by humans are widespread across the tree of life. Following human arrival, many reptile and amphibian species went extinct in New Zealand (*15*). At least 581 prehistoric extinctions have been documented in birds (*16*). Nevertheless these numbers may still be lower than the true number (*17*) if undocumented “dark extinctions” (*18*) or “dark extirpations” are widespread. Early humans contributed to extinctions in a variety of ways including: hunting (*19*), introductions of non-native species, and alteration of natural habitats (forested areas often became heavily degraded and more open) (*20-23*).

Here, we assess whether early humans have caused sensitive species to be lost leading to communities that are less sensitive to habitat modification. To accomplish this we use estimates of human populations dating back >12,000 YBP (*24*) and pair this with modern-day bird community data (*25*). Total human population size serves as an integrated index of all human activities, inclusive of land-use change, direct hunting pressure, and indirect effects from human commensals. We aim to differentiate between two non-mutually exclusive hypotheses, the extinction filter hypothesis and the adaptive resilience hypothesis. The extinction filter hypothesis predicts that natural habitats in regions with large historical human presences have suffered dark extinctions and lost their most sensitive species, such that in the present day they will have lower diversity in comparison to similar regions that had lower historical human presence. Because species occurring in human-modified habitats are by definition at least moderately tolerant of human impacts and not susceptible to the extinction filter, this hypothesis further predicts that similar types of human-modified habitats across regions should have similar levels of species richness, regardless of ancient human presence. In contrast, the adaptive resilience hypothesis predicts that sustained human presence should generate an increase in tolerant species (either through adaptation, acclimation, or selective immigration). As such, this hypothesis predicts an increase in human-associated species in areas with sustained human presence. Such a scenario may be possible as a wide variety of animals have been shown to adapt to human influences (*26-28*), and human commensals are frequently among the most successful invasive species, demonstrating an ability to colonize human impacted areas through time (*29, 30*).

### Historical human influence on community sensitivity

We used the PREDICTS Database (*25*) to obtain contemporary bird community data in different land uses across the globe. We focused our analysis on three major categories of land-use (see supplemental description of classification procedure). Primary vegetation represents the undisturbed natural habitat for the ecoregion in question. We represent modern human-dominated land in two alternative forms: structurally “complex” and “simple” agriculture. Complex agriculture contains large bushy or tree like crops that are generally more than 2 meters tall. These crops may offer more vegetative structure for organisms to use as habitat, partially mimicking some forms of primary vegetation. Such forms of agriculture include coffee, nut, and tree plantations or orchards. In contrast “simple agriculture” has lower stature crops that grow less than 2m tall such that they have relatively little useable habitat structure. “Simple agriculture” includes pasturelands in addition to low stature row crops. To best exploit the paired nature of the data, we removed all studies which did not contain primary vegetation, and at least one other land use category.

We used 54 different studies, with 3,975 unique sites, 55,267 species-by-site pairings, and 2,645 total bird species (roughly one fourth of all named species).Then we matched these biodiversity surveys with human population density data and land use estimates obtained from the History Database of the Global Environment (*24*) at a 5 arc minute resolution for multiple time periods between 12,000 YBP to 2000 CE. In our analyses we controlled for mean annual temperature and precipitation at 1km resolution (*31*), to ensure that climate related differences across the globe were not spuriously driving observed trends. Controlling for the independent effects of climate, overall bird species richness was greatest in primary vegetation and lowest in simple agriculture. However, human population density throughout time had powerful influences on species richness patterns. Human population density has increased greatly in the last 12,000 years, population densities during the oldest time period 12,000 YBP (0-2.1 humans per km^2^ at sites) were very low compared to the most recent time period 2000 CE (0-1,800 humans per km^2^). Of all time periods considered, the oldest, 12,000 YBP was the best at describing the data (Table 1), suggesting that extremely early human impacts have compounded to structure modern day bird communities (human density by land use interaction effect: negative binomial mixed effect model, LRT = 47.7, p < 0.001). Areas with greater historical human population sizes have less sensitive communities (Figure 2). We find support for the dark extinction hypothesis, that ancient humans have increased the resilience of communities by eliminating species that would otherwise occur in primary vegetation. Specifically, fewer species persist in today’s primary vegetation in areas where human populations were high 12,000 year ago, while the number of species occurring in structurally simple agriculture is relatively unimpacted by the presence of ancient human populations (Figure 2). Within primary vegetation, predicted species richness declines by 20% when comparing areas with no humans 12,000 YBP to areas with relatively high human populations (0.67 humans per km^2^, the 95% quantile of surveyed bird communities analyzed). This level of species richness loss within primary vegetation is similar to the amount of change between primary vegetation and agriculture in areas which lacked humans 12,000 YBP, where average bird species richness declines by 22% from primary vegetation to structurally simple agriculture, and 19% from primary vegetation to complex agriculture. In contrast, in areas with high pre-historic human populations (0.67 humans per km^2^), modern day species richness is sufficiently low in primary vegetation that species richness is not significantly different between landcovers, with predicted declines of only 2% in simple agriculture, and 11% in complex agriculture.

**Fig. 1.**
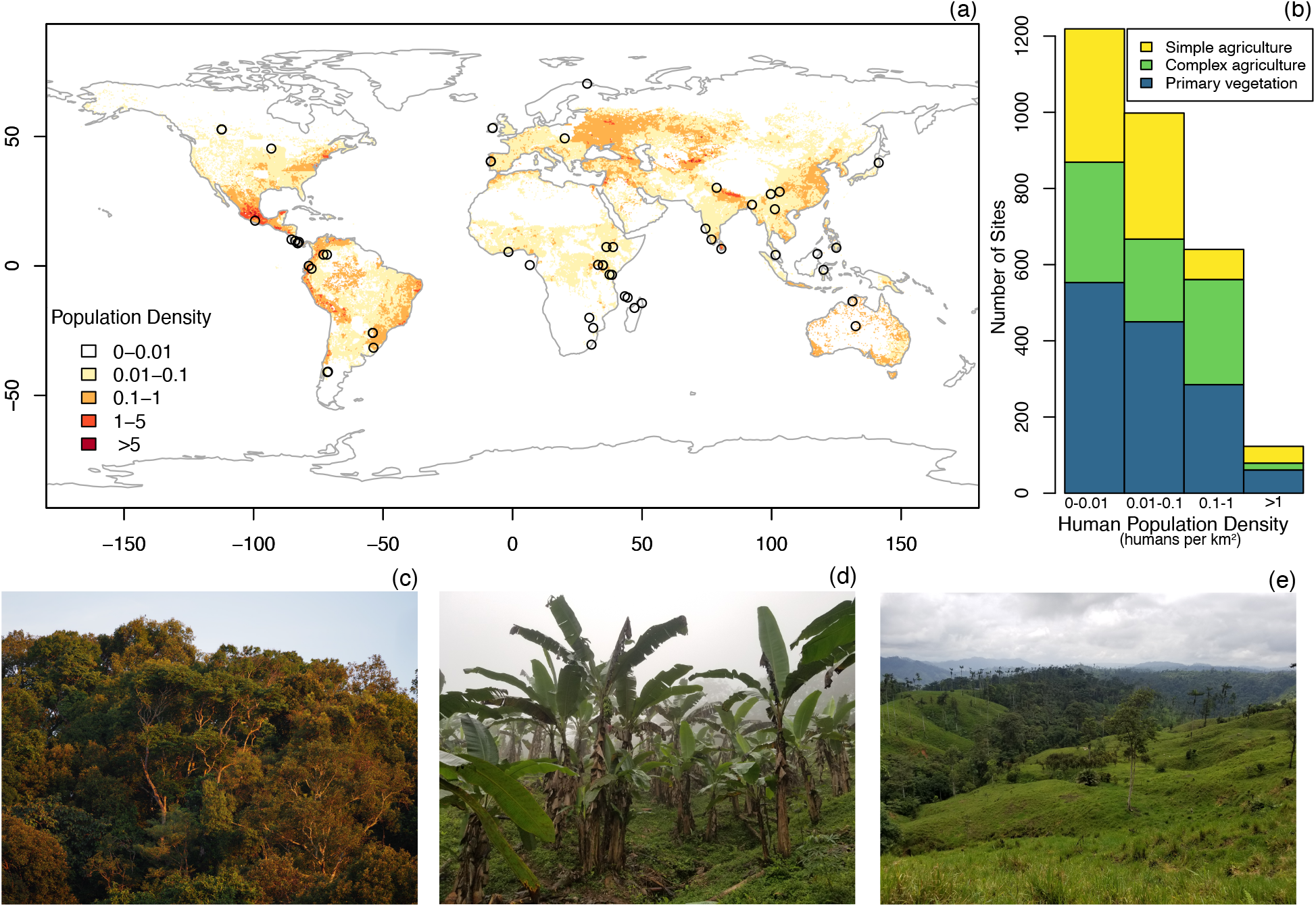
**A)** Map displaying human population density (humans per km^2^) estimates from Hyde 3.2 for 12,000 YBP (Years Before Present). Points represent the central location of each of the 54 studies included in our analysis. B). Number of sites falling within the population density classes shown on the map, colored by land use classification, C) primary vegetation, D) complex agriculture, and E) simple agriculture.

**Table 1.** Results from models testing the impacts of human population density on species richness at different time periods, all models contain the full set of data from 54 studies. LU indicates primary land-use of the survey location in the present day, MAT is mean annual temperature (bio1), whereas MAP is mean annual precipitation (bio12), both from the WorldClim dataset. Asterisks indicate significance level of the variables or interaction effects in question, based on a likelihood ratio test. *P-value >0*.*05 NS, <0*.*05 *, <0*.*01**, <0*.*001\*\*\**.

| Model # | Time period | AIC | ΔAIC | Human population × LU | MAT × LU | MAP |
| --- | --- | --- | --- | --- | --- | --- |
| 1 | 12,000 YBP | 23341.3 | 0 | *** | *** | *NS* |
| 2 | 4,000 YBP | 23382.0 | 40.7 | ** | *** | *NS* |
| 3 | 2,000 YBP | 23390.7 | 49.4 | * | *** | *NS* |
| 4 | 500 CE | 23388.2 | 46.9 | * | *** | *NS* |
| 5 | 1000 CE | 23379.9 | 38.6 | ** | *** | *NS* |
| 6 | 1500 CE | 23384.9 | 43.6 | * | *** | *NS* |
| 7 | 1700 CE | 23386.3 | 45 | * | *** | *NS* |
| 8 | 1800 CE | 23389.0 | 47.7 | * | *** | *NS* |
| 9 | 1900 CE | 23381.4 | 40.1 | *** | *** | *NS* |
| 10 | 2000 CE | 23382.4 | 41.1 | *** | *** | *NS* |

**Fig. 2.**
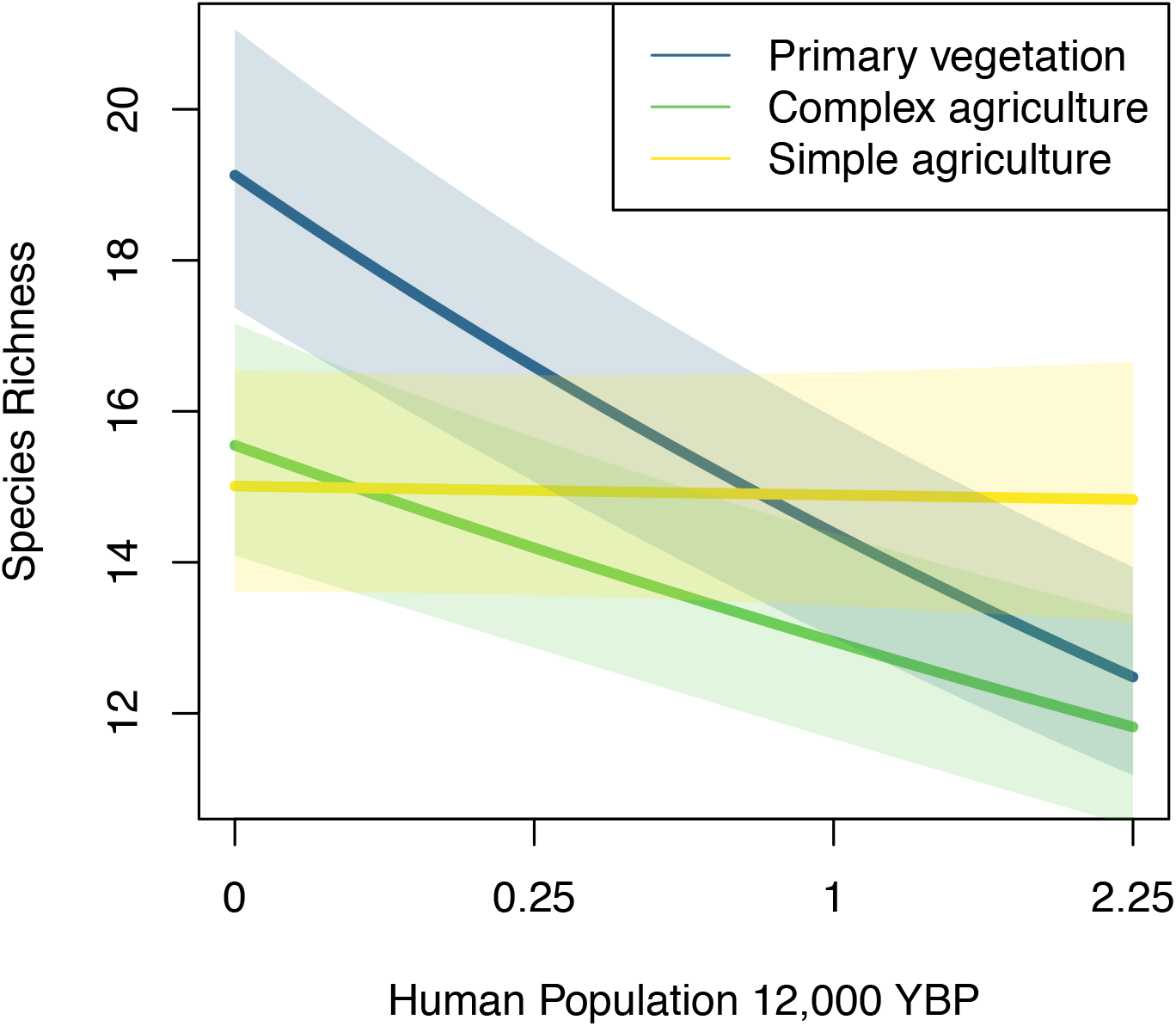
Model predicted relationships between human population density ~12,000 YBP (humans per km^2^) and species richness, shaded areas represent model-predicted standard error.

In order to understand the mechanism behind species loss in primary vegetation we assessed whether such declines reflect a loss of primary vegetation specialist species, as opposed to loss of species capable of inhabiting both disturbed and natural habitats. We classified species as primary vegetation specialists if they were only ever found within primary vegetation within the study in question. We then tested whether primary vegetation communities, with larger human populations 12,000 YBP, had fewer primary vegetation specialists, as opposed to multi-habitat generalists. We found only modest support for this hypothesis: primary vegetation specialists tended to account for a smaller number of species occurring in primary vegetation where human populations were large 12,000 YBP, however this result is only marginally significant (p-value = 0.07), with a predicted 8% decrease in the proportion of primary vegetation specialists at sites in areas with zero humans 12,000 YBP compared to areas with a population density of 1 humans/km^2^. This suggests that ancient human impacts extend beyond just the most specialized species, and affected habitat generalists as well.

### Early human impacts on phylogenetic diversity

Past research has suggested that sensitivity to habitat modification and extinctions are not randomly distributed throughout the tree of life (*32*). Evolutionary distinct species, those with few extant close relatives, are believed to amongst those most frequently lost when natural habitats are converted to those for human use (*33, 34*). We assessed the phylogenetic diversity, phylogenetic clustering, and median (and mean) evolutionary distinctiveness of communities to assess whether historical human occupancy has disproportionately reduced phylogenetic diversity by eliminating evolutionarily unique species. Using a phylogenetic tree of all birds (*35*), we first tested if phylogenetic diversity decreases in areas with higher human populations 12,000 YBP, while controlling for the effects of climate and study. Similar to trends with species richness, phylogenetic diversity decreases in primary vegetation and complex agriculture while in simple agriculture remains relatively unchanged by human population size (human density by land use interaction effect: linear mixed effect model Chi^2^ = 15.5, p = 0.004; Figure 3a). Since phylogenetic diversity is tightly correlated with species richness, we next assessed phylogenetic clustering (mean phylogenetic distance) within communities, to determine if clustering increases in communities with long histories of human occupation, as would be expected if evolutionarily distinct species are disproportionately eliminated. In areas with minimal human influence in the deep past, communities across land use types showed roughly equivalent degrees of phylogenetic clustering. However, contrary to our expectation, as historical human influence increased, communities in primary vegetation and complex agriculture both became less clustered suggesting that the species that remain are more broadly sampling the avian tree of life (human density by land use interaction effect: Chi^2^ = 15.6, p =0.004; Figure 3b). If extinctions occurred, we might expect to see an increase in the evolutionary distinctiveness (ED) (*36*) in bird communities in areas where human populations were high in ancient times, as extinctions of closely related species leave remaining extant species more isolated on tree of life. Our results support this hypothesis, as median (and mean) ED values were similar across habitats with low human populations in ancient times. However, in areas with high human populations 12,000 YBP, the mean evolutionary distinctiveness of today’s communities is higher in primary vegetation, whereas in simple agriculture it is relatively unchanged (human density by land use interaction effect: Chi^2^ = 21.3, p <0.001; Figure 3c). These results suggest that the species lost from primary vegetation in areas long occupied by humans represent species that were closely related to other species that still occur at the sites, and come primarily from recently radiating clades from across the globe.

**Fig 3.**
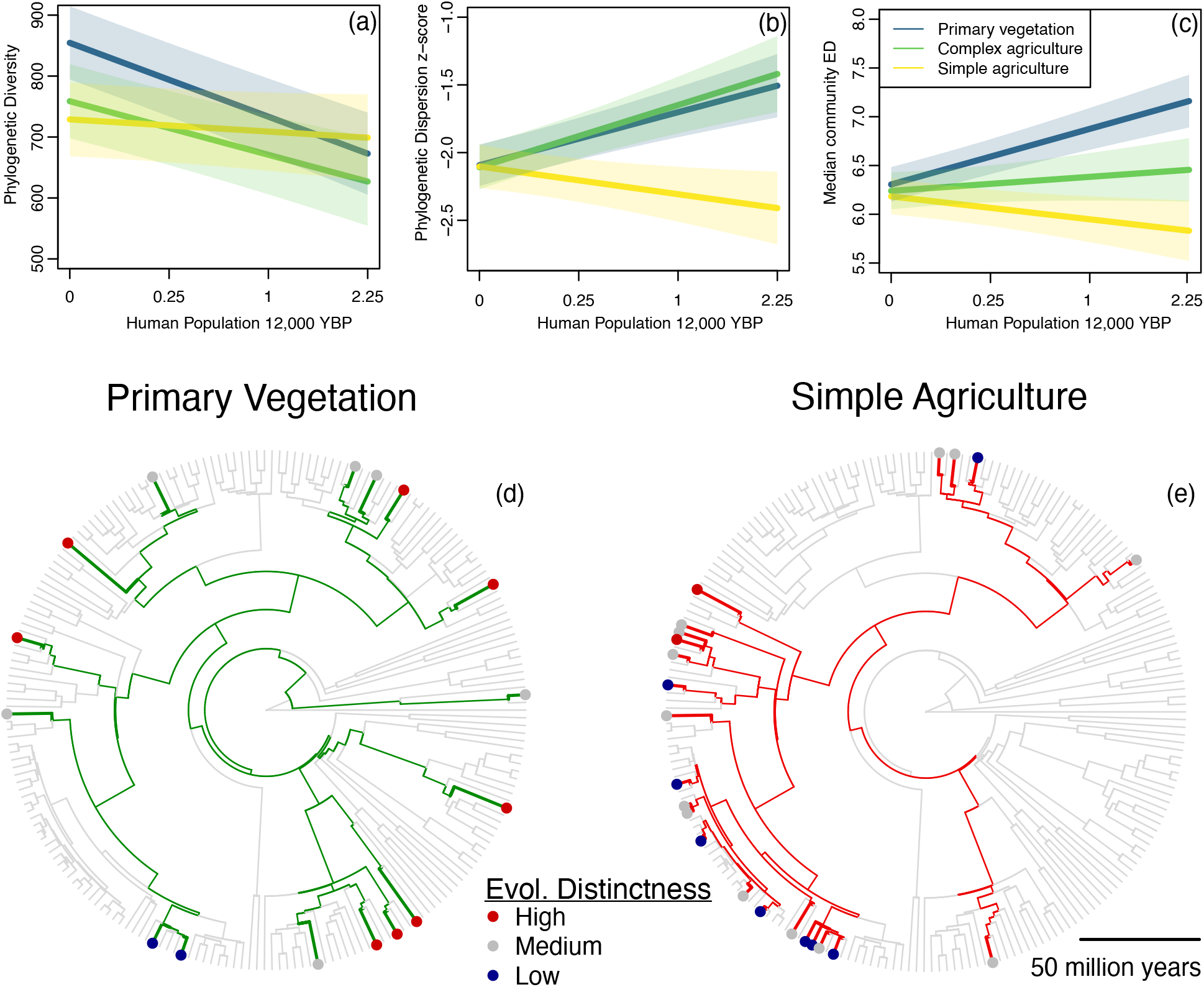
**A**) Model predicted phylogenetic diversity (Faith’s PD). B) Model predicted mean phylogenetic distance (MPD) Z-score with human population density 12,000 YBP (humans per km^2^) C) Model predicted median evolutionary distinctiveness (ED) relationship with human population density 12,000YBP. D) Phylogenetic tree of species found in the western Andes (50) where the human population density 12,000 YBP was relatively high (>1 humans per km^2)^, at a site in primary vegetation. E) Bird species found at a simple agriculture site in the same study, note the greater clustering in simple agriculture and they greater number of low ED species.

Taken together our results suggest that impacts from early humans extend far beyond megafaunal extinction. Birds have the greatest amount of available data out of vertebrate groups, however we believe this to be a phenomena that is likely widespread across other taxonomic groups. In particular less mobile organisms (e.g. reptiles and amphibians) may bear even stronger signals of direct human presence (*37, 38*), since their ability to recolonize areas are hindered by their low mobility (*39, 40*). Indeed, plant diversity in areas previously occupied by humans often still reflects this history of anthropogenic impacts even thousands years later (*38*). The data suggest that humans have acted as an extinction filter, and that even early human populations are linked to declines in bird species richness in primary vegetation. While we find a negative relationship between human population density and avian biodiversity, both today and in the past, historical patterns of human occupancy up to 12,000 years ago better predict modern day patterns of bird diversity across land uses than do more recent population densities. The greater predictive power of the earliest populations distributions occurs despite the low population densities at this time. We hypothesize that even at relatively low absolute densities, human impacts on landcover and through hunting, compounded over thousands of years, resulted in sustained species losses in the landscapes involved.

### Global vs. local extinctions

The declines in species richness we document in primary vegetation could manifest as a result of either merely local or global extinction of the species at hand. Evolutionary distinctiveness hints that we have perhaps underestimated the extent to which global extinctions occurred dating back in time, and thus have pruned back bushier branches of the tree of life, while counter intuitively leaving longer branches relatively less affected. The exact mechanism by which the extirpations we infer in primary vegetation occurred is not certain, as hunting, human-driven fire regimes, and land use change often all occurred together. However, habitat modification may have been the primary way in which species were impacted by early humans, as communities in primary vegetation and complex agriculture show similar patterns of decreasing species richness and phylogenetic diversity, whereas in simple agriculture the community is largely unimpacted by historical human presence. Complex agriculture and primary vegetation have structural similarities, particularly in forested regions, in that they tend to have at least some degree of canopy cover and greater vegetative structure. If prehistoric humans were primarily impacting birds through habitat modification, and this led forest to be converted to more open and less structurally complex habitat then species which were dependent on forest/closed canopy might decline, while species which are tolerant of more open areas may have been left relatively unimpacted or increased due to greater availability of suitable habitat. In contrast, if hunting were the primary mechanism by which species were being impacted by humans, then we might expect to see equal differences across land use types. But regardless of mechanism, communities in primary vegetation bear the brunt of the impacts, which runs counter to the idea that intact primary vegetation constitute areas untouched by human influence and reinforces the significant role of humans in natural landscapes across the globe (*41*). If the differences in species richness reported here do not represent global extinctions, but rather are instead merely local extirpations, then it suggests that these species have not been able to recolonize primary vegetation since. A potential mechanism may be that remnant patches of primary vegetation are too small or too isolated for successful colonization and persistence, such that areas of long human habitation have more fully paid the extinction debt incurred by initial human impacts (*42, 43*). Without sub-fossils or fortuitously preserved specimens, determining whether these less diverse areas are a result of true extinctions is difficult. The alternative hypothesis, that adaptation or acclimation is responsible for a reduced sensitivity of communities to disturbance is not supported by our study: there was no indication of a positive relationship between species richness in disturbed habitats and pre-historic human population size.

### Implications for conservation

Growing recognition of the severity of human impacts on the environment has led to a movement to rename our current geological epoch “the Anthropocene”. The exact start of such an epoch has been debated, with most suggesting a relatively recent start date in the mid 1900s for geological dating purposes (*44-46*). Regardless of the official start date, our results suggest that human impacts dating back to 12,000 YBP are widespread and pervasive.

Underestimating the import of such early impacts may hamper our understanding of biodiversity patterns and conservation needs. Contemporary patterns of biodiversity and community structure are considered to be a product of historical factors and contemporary conditions (*47*), however historical human presence is rarely considered as a potential driver of differences in diversity. Including influences of early humans in studies may help resolve what generates different biodiversity amounts in regions with similar climates and environments (*47*). Our results shed some light on where dark extinctions likely occurred over thousands of years of burgeoning human populations. As evidence emerges that combinations of past and current factors together drive community sensitivity to human-impacts, this diversity of drivers needs to be integrated into a unified framework. Doing so will enable efficient use of the limited resources available for conservation, and enhance our ability to identify areas most in need of preservation based on conservation value and the sensitivity of the communities within.

## Acknowledgments

We thank Edita Folfas, Zachary Lange, and Mara Pineau for feedback on drafts of this paper. We also thank all researchers who have done the important work to collect the data used in this study and make it publicly available for research.

## Funding

The Frishkoff lab was supported by NSF Award #2055486.

## Author contributions

Conceptualization: AHM

Methodology: AHM, LOF

Formal analysis: AHM

Visualization: AHM, LOF

Supervision: LOF

Writing – original draft: AHM

Writing – review and editing: AHM, LOF

## Competing interests

Authors declare that they have no competing interests.

## Data and materials availability

All data were from publicly available sources. Bird community data are from the PREDICTS database at https://www.nhm.ac.uk/our-science/research/projects/predicts.html. The climate data used are from the WorldClim data set and are available at https://www.worldclim.org/data/index.html. Data on human population density are from the HYDE dataset and are available at https://landuse.sites.uu.nl/datasets/. The phylogenetic tree used for phylogenetic analyses is available at https://vertlife.org/data/. The final data set used for analyses and code to run models are available at https://doi.org/10.6084/m9.figshare.25021310.v1.

## Supplementary Materials

Materials and Methods

Figs. S1

Tables S1

References (*48-49*)

## Materials and Methods

### Trends in species richness

We obtained community data from the PREDICTS database and combined the original data release with the 2022 additional data release. We filtered our dataset to only contain birds, as they have the largest dataset amongst vertebrate species with the most widespread distribution of sites. A total of 69,7801 of species by site pairings existed before we cleaned the dataset. We removed studies which contain small taxonomic scope such as single species studies, single land uses, or no primary vegetation. Within studies we eliminated sites which had unequal sampling effort. After cleaning records, we retained 54 different studies, with 3,975 unique sites, 55,267 species-by-site pairings, and 2,645 total bird species. We classified land use into 5 different categories based off authors descriptions. Three of these are forms of natural land: Primary vegetation representing the undisturbed habitat for the ecoregion in question, mature secondary vegetation (generally representing vegetation that has regrown for at least 50 years since clearing) and young secondary vegetation (less than 50 years since clearing). We represent human-dominated land in two alternative forms: structurally “complex” and “simple” agriculture. The greatest number of sites were primary vegetation 1,559, followed by complex agriculture 894, simple agriculture 860, young secondary vegetation 649, and mature secondary vegetation 13. We present results in main text for primary vegetation, complex agriculture, and simple agriculture as the interest is in how species richness is impacted by land-use change, however patterns found in simple agriculture and young vegetation are similar (Figure S1). We tested the influence of human dominated land uses on species richness by including an interaction between land use type and human population density as measured in different time periods (10 total time periods from 12,000 YBP to 2000CE), to determine if differences in species richness between land covers is altered by the history of human presence. Because human population densities are bounded by zero, a large proportion of the data were small values, and relatively few had large values. To reduce the leverage of large values we therefore applied a square root transformation of human population density before analysis, though results are broadly concordant regardless of whether square root or raw data are used (Table S1). We controlled for the effects of climate by including mean annual precipitation, and an interaction between land use and mean annual temperature (because effects of habitat modification can be stronger in warmer areas) as fixed effects, and study as a random effect using generalized linear mixed effects models with a negative binomial distribution in the *glmmTMB* in R (*48*).

### Specialization

We classified species as primary vegetation specialists if they were only ever found within primary vegetation within the study in question. As such, the same species could be classified as primary vegetation specialists in one study and not in another, because habitat use can vary across species ranges. We tested whether primary vegetation communities where there were larger human populations 12,000 YBP had fewer primary vegetation specialists, as opposed to multi-habitat generalists (everything that is not a primary vegetation specialist). We filtered our dataset down to only include primary vegetation sites, a total of 1,559 sites from 54 studies. We used generalized linear mixed effects models with a binomial distribution in the *glmmTMB* in R (*48*) with the number of primary specialists observed at a site compared to generalist species at a site as the response variable. We included human population size 12,000 YBP, mean annual temperature, and mean annual precipitation as fixed effects, and controlled for study as a random effect.

### Phylogenetic diversity

We obtained the Ericson-backbone tree of all birds (*35*), and calculated Faith’s phylogenetic diversity for each site using the *pd* function in Picante (*49*). We tested if phylogenetic diversity decreases in areas with higher human populations 12,000 YBP, by using phylogenetic diversity as our response variable and including interactions between human population size 12,000 YBP and land use, mean annual precipitation, and an interaction between land use and mean annual temperature as fixed effects, while controlling for the effect of study as a random effect using generalized linear mixed effects models with a gaussian distribution in the *glmmTMB* in R (*48*).

We tested how clustered our communities were by calculating phylogenetic dispersion. To do so we calculated mean pairwise distance within sites and z-scores from comparisons to a tip-swap null distribution (1,000 simulations) using the *ses*.*mpd* function in Picante (*49*). We extracted the z-scores and then asked what controlled whether a community was phylogenetically more clustered, or more over dispersed in areas with large prehistoric populations. We used z-scores as our response variable and including interactions between human population size 12,000 YBP and land use, mean annual precipitation, and an interaction between land use and mean annual temperature as fixed effects, while controlling for the effect of study as a random effect using generalized linear mixed effects models with a gaussian distribution in the *glmmTMB* in R (*48*). We assessed mean and median evolutionary distinctiveness (ED) of bird species within sites by obtaining values of ED from (*36*). We used median ED (and mean in separate models) as response variables and including interactions between human population size 12,000 YBP and land use, mean annual precipitation, and an interaction between land use and mean annual temperature as fixed effects, while controlling for the effect of study as a random effect using generalized linear mixed effects models with a gaussian distribution in the *glmmTMB* in R (*48*).

## Notes

### Competing Interest Statement

The authors have declared no competing interest.

