## Supplemental Materials for "Ancient occupation by humans leads to missing bird diversity in otherwise natural habitats"

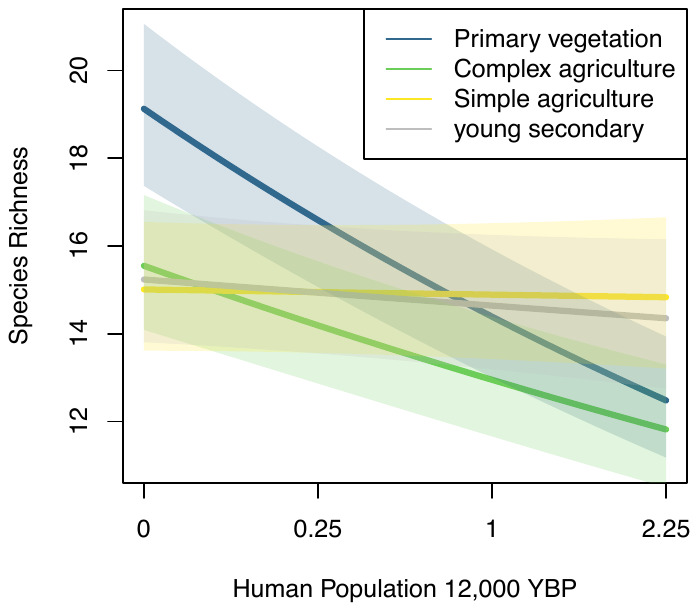

Fig. S1. Model predicted relationships between human population density ~12,000 YBP (humans per km^2^) and species richness, shaded areas represent standard error.

**Table S1.** Results from models testing the impacts of human population density on species richness at different time periods, all models contain the full set of data from 54 studies*.* This set of models is based off raw values of Human population density, not the square root as presented in the main text. LU indicates primary land-use of the survey location in the present day, MAT is mean annual temperature (bio1), whereas MAP is mean annual precipitation (bio12), both from the WorldClim dataset. Asterisks indicate significance level of the variables or interaction effects in question, based on a likelihood ratio test*. P-value >0.05 NS, <0.05 *, <0.01**, <0.001***.*

Type or paste caption here. Create a page break and paste in the table above the caption.

| **Model #** | **Time period** | **AIC** | **ΔAIC** | **Human Population * LU** | **MAT * PLU** | **MAP** |
| --- | --- | --- | --- | --- | --- | --- |
| 1 | 12,000 YBP | 23331 | 0 | *** | *** | NS |
| 2 | 4,000 YBP | 23388.7 | 57.7 | . | *** | NS |
| 3 | 2,000 YBP | 23392.4 | 61.4 | * | *** | NS |
| 4 | 500 CE | 23391.4 | 60.4 | * | *** | NS |
| 5 | 1000 CE | 23390.8 | 59.8 | * | *** | NS |
| 6 | 1500 CE | 23393.2 | 62.2 | NS | *** | NS |
| 7 | 1700 CE | 23392.4 | 61.4 | * | *** | NS |
| 8 | 1800 CE | 23393.5 | 62.5 | * | *** | NS |
| 9 | 1900 CE | 23389.4 | 58.4 | ** | *** | NS |
| 10 | 2000 CE | 23386.4 | 55.4 | ** | *** | NS |
